# Adaptive zinc tolerance is supported by extensive gene multiplication and differences in cis-regulation of a CDF transporter in an ectomycorrhizal fungus

**DOI:** 10.1101/817676

**Authors:** Joske Ruytinx, Laura Coninx, Michiel Op De Beeck, Natascha Arnauts, François Rineau, Jan V. Colpaert

## Abstract

Abiotic changes due to anthropogenic activities affect selection regimes for organisms. How trees and their mycorrhizal symbionts adapt to altered environments in heterogeneous landscapes is of great interest. With a global distribution and multiple adaptive phenotypes available, *Suillus luteus* is an excellent ectomycorrhizal model to study evolutionary dynamics of local adaptation. We assessed pathways of homeostasis and detoxification in *S. luteus* isolates, displaying contrasting Zn tolerance phenotypes to identify mechanisms underlying adaptive Zn tolerance. Using 30 randomly selected isolates sampled at metal contaminated and control sites, we documented Zn tolerance phenotypes, assessed the link with identified candidate genes and explored its genetic basis via targeted amplicon sequencing and qPCR. Zn tolerance phenotypes covering a continuum from Zn sensitive to hypertolerant were identified and inversely correlate with cellular Zn accumulation. Gene expression of *SlZnT2*, encoding a putative Zn transporter explains 72% of the observed phenotypic variation. *SlZnT2* copy number varies among isolates and different promotor genotypes were identified. Rapid adaptation in this species is supported by the cumulative effect of gene copy number variation and differences in cis-regulation and might be triggered by environmental stress rather than being the result of standing variation.

**Originality - significance statement:** To the best of our knowledge, this is the first study linking genotypes to adaptive phenotypes in mycorrhizal fungi. It is unique in the way it combines evolutionary and functional genetics to allow a significant advance in the understanding of responses to environmental stress factors in general and, to soil metal pollution in particular. A better understanding of adaptive tolerance mechanisms in keystone symbiotic fungi is paramount for developing impactful phyto and mycoremediation strategies for metal polluted waste land and to predict the impact of future environmental change on mycorrhizal diversity and ecosystem functioning.

## Introduction

Human-induced environmental changes, including pollution, soil erosion and climate change, and their impact on ecosystem functioning are of major concern. Plant and associated microbial communities are intensively studied to predict how abiotic factors affect nutrient cycling and species diversity (e.g. Bradford et al., 2019; Franklin et al., 2016; Fernandez et al., 2016, Kyaschenko et al., 2017). Climate parameters and edaphic factors influence relative abundances and species composition of plant and microbial communities (Castaño et al., 2018; Op De Beeck et al., 2015; Wang et al., 2018). Hostile environments challenge species and survival requires contemporary local adaptation. Historically, mechanisms of local adaptation in plant species, including physiology and genetics, attract the interest of many scientists (recent examples include Selby & Willis, 2018; Ferrero-Serrano & Assman, 2019). However, for many plant species and in particular long living trees, microbial mutualists including ectomycorrhizal fungi largely determine their environmental plasticity, adaptive and invasive potential (Wardle et al., 2004; Hayward et al., 2015; Dickie et al., 2017; Policelli et al., 2019). Despite growing interest, it remains unclear how abiotic factors act on genetic and functional diversity of mycorrhizal symbionts. Examples of local adaptation in mycorrhizal fungi are limited and related to climate parameters, nutrient availability or presence of potentially toxic trace metals (Colpaert et al., 2004; Johnson et al., 2010; Jourand et al., 2010; Zhang et al., 2019). Yet, mycorrhizal fungi showing local adaptation protect their host plant effectively from the environmental challenge posed, resulting in a balanced nutrient supply despite unfavourable availabilities (Johnson et al., 2014; Adriaensen et al., 2005; Krznaric et al., 2009; Jourand et al., 2014). An interesting case are Suilloid fungi, with multiple pioneer species within the genera *Suillus* and *Rhizopogon* showing environmental adaptation towards high metal availability in soils and, independently evolved Zn, Cd and Cu tolerance phenotypes (Colpaert et al., 2004; Colpaert et al., 2011). Adaptive phenotypes of these species presumably support the establishment of pines on severely contaminated soils (Colpaert et al., 2011). Evolutionary dynamics of local adaptation and the genomic features governing the adaptive potential of these particular Suilloid species are as for other ectomycorrhizal fungi not yet explored despite their putative importance for the understanding of pine establishment on waste lands.

Rapid adaptation to strong human induced shifts in the environment is repeatedly linked to discrete traits. Discrete traits are traits with only a few possible phenotypes, clearly falling in discrete classes. Well known examples include shifts in populations from white to black-coloured peppered moths due to industrialisation (van’t Hof et al., 2016) and from non-resistant to insecticide resistant Drosophila (Daborn et al., 2002). In this case, the phenotype is controlled by one (or few) genetic locus of major effect and environmental shifts result in frequency shifts of the alleles at that particular locus allowing rapid adaptation of the population. A strong selective sweep or dramatic change in allele frequency was detected among ectomycorrhizal *Suillus brevipes* populations sampled in North-American coastal and mountainous regions and considered as a genomic signature of environmental adaptation. The highly differentiated genetic locus contains a transporter gene with putative function in adaptive salt tolerance (Branco et al., 2015). In the same species, genetic differentiation across the North-American continent correlates with climate parameters (Branco et al., 2017). Phenotypic differentiation among *S. brevipes* populations ultimately resulting in an improved environmental fitness was not assessed as a proof of true local adaptation. Neither were functions of the identified candidate genes.

Evidence of rapid environmental adaptation thanks to differentiation in quantitative traits, i.e. traits of which the phenotypes can be expressed as quantitative values covering a continuum rather than being attributed to discrete classes, is only emerging recently. Examples include adaptation of yeast towards toxins and of American house mouse towards latitude (Naranjo et al., 2015; Phifer-Rixey et al., 2018). A cumulative action of different genetic loci or eQTLs (expression quantitative loci) on the expression of a particular gene associates with the adaptive phenotype in these examples (Naranjo et al., 2015; Mack et al., 2018). In general, genetic variants affecting gene expression are a major source of phenotypic variation and allow modulation of existing cellular pathways. However, individual (e)QTLs are difficult to identify when they have a small effect size (contribution to the final phenotype is low) and/or the cohort of individuals under investigation is small (Anderson et al., 2011). In fungi, genetic loci were associated with complex traits such as virulence using <25 individuals. Identified loci however, all displayed a strong effect on phenotype (Taylor et al., 2017).

*Suillus luteus* is a soil-borne fungal species associating with pine trees in ectomycorrhizal symbiosis. In particular, in primary successions or colonisation of disturbed sites this species is determining the success of pine establishment across native and non-native ranges (Hayward et al., 2015; Policelli et al., 2019). Adaptive tolerance towards high concentrations of Zn was described for populations thriving on severely heavy metal contaminated soils (Colpaert et al., 2004). The gene repertoire for all kinds of metal detoxification pathways is present in ectomycorrhizal fungi (Bolchi *et al.*, 2011). In *S. luteus*, different Zn transporters with a function in cellular Zn uptake and distribution were characterized (Coninx et al., 2017; Ruytinx et al., 2017) and genes encoding additional proteins putatively involved in homeostasis, detoxification and damage repair were identified (Muller et al., 2007, Ruytinx et al., 2011, Nguyen et al., 2017). In the current study, we hypothesize contrasting Zn tolerance phenotypes in *S. luteus* to be due to alterations in general mechanisms of Zn homeostasis, detoxification or damage repair. The differential use of these mechanisms among isolates is explored by specific enzyme and gene expression assays. We document natural Zn tolerance phenotypes and aim at identifying the physiological, molecular and genetic basis of Zn tolerance using a targeted approach. An improved understanding of physiological and molecular mechanisms underlying adaptive phenotypes in mycorrhizal fungi, including their genetic basis and evolutionary dynamics could assist in the sustainable management of infertile agricultural soils and restoration of abandoned or polluted industrial land. Even so, it would ultimately allow to model and predict the impact of future abiotic change on fungal genetic and functional diversity.

## Results

### Enzyme activity and lipid peroxidation in contrasting phenotypes

The enzyme activity of two groups of anti-oxidative enzymes, i.e. catalases (CAT) and superoxide dismutases (SOD) was measured in *S. luteus* isolates showing contrasting Zn tolerance phenotypes. A significant decrease in CAT activity after exposure to 1000 μM Zn was detected in two isolates despite their contrasting Zn tolerance phenotypes (Lm8 and Mm7; fig. 1a). SOD activity was effected upon Zn treatment for all Zn-sensitive *S. luteus* isolates. However, the Zn concentration triggering the effect differed among isolates (fig. 1b). Thiobarbituric acid reactive substances (TBARS) were measured as a product of damage produced by oxidative stress. All isolates show a comparable level of malondialdehyde (MDA) production and for none of them an effect of Zn exposure on MDA production could be detected (fig. 1c).

**Figure 1.**
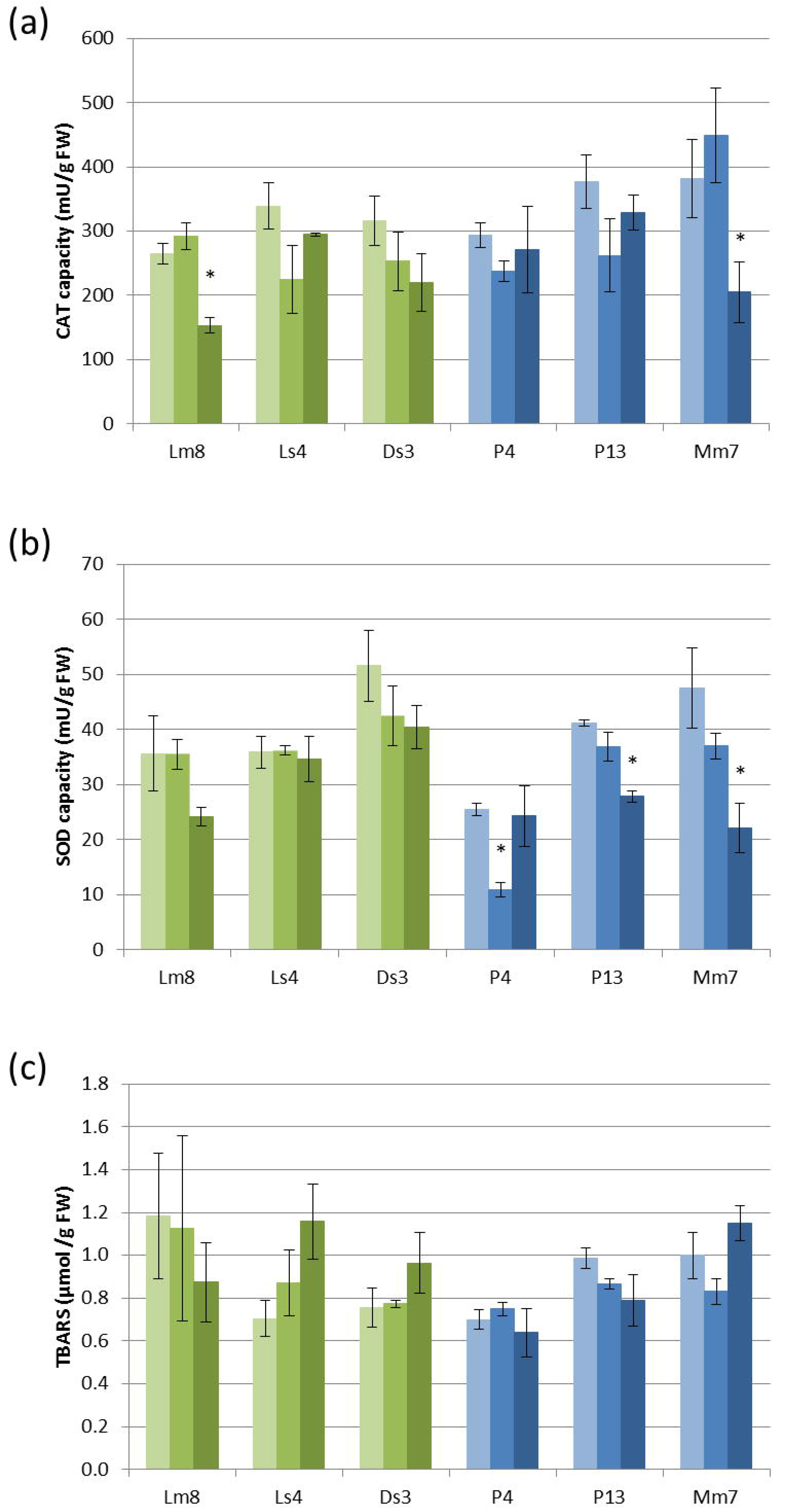
Enzyme activities and lipid peroxidation measured in individual Zn tolerant (green) and Zn sensitive (blue) *S. luteus* isolates upon 48h exposure to different Zn (20, 200, 1000 μM) concentrations. (a) catalase activity; (b) superoxide dismutase activity; (c) lipid peroxidation measured as TBARS (thiobarbituric acid reactive substances). Higher Zn concentrations correspond to darker bars. Data are the average of 3 replicates +/− S.E.; significant differences (p<0.05) compared to the control (20μM) are indicated by *.

### Transcription of selected candidate genes in contrasting phenotypes

On the transcript level, several genes encoding proteins involved in Zn homeostasis and metal detoxification were measured upon exposure to different concentrations of external Zn (fig. 2a-h). A Zn importer (ZRT1), a tonoplast localized transporter involved in Zn accumulation (ZnT1) and a putative Zn transporter of unknown localization (ZnT2) were studied. Transcripts of *ZRT1* decline with increasing external Zn concentration in all isolates. In general, transcript level of this transporter is higher in isolates with a more sensitive phenotype (2-10x) compared to tolerant phenotypes when considering the same external Zn concentration (fig. 2a). In contrast, ZnT1 transcript level is comparable in all isolates and not responsive to external Zn concentration in most of the isolates (except 2 Zn sensitive isolates at 1000 μM Zn; fig. 2b). Biased transcript levels between isolates of both Zn tolerance phenotypes, with fold changes ranging from 5-100x, were observed for ZnT2 (fig. 2c). This gene is constitutively highly expressed in more tolerant isolates whereas the transcript is almost absent in more sensitive isolates.

**Figure 2.**
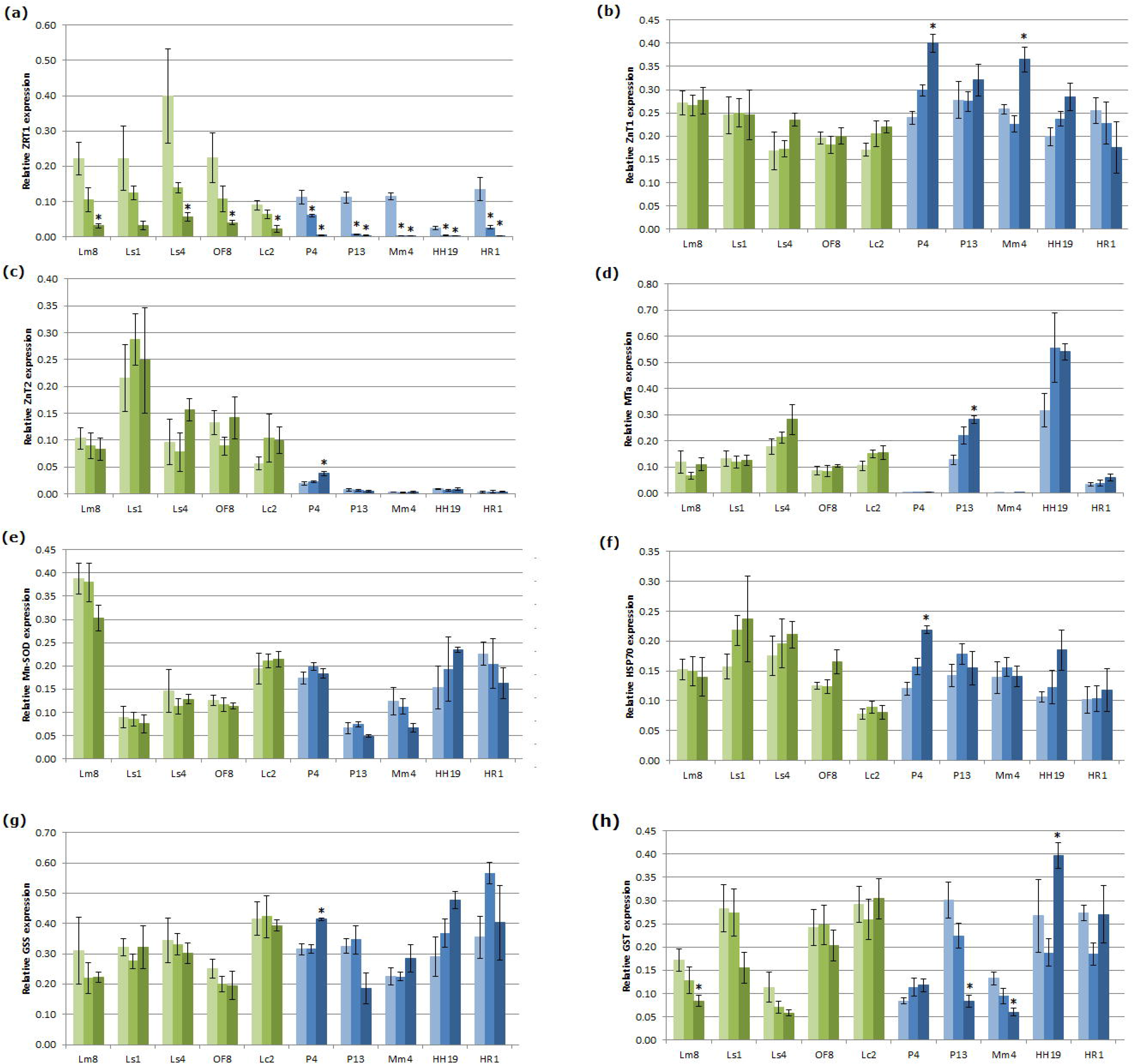
Relative expression of selected genes measured in individual Zn tolerant (green) and Zn sensitive (blue) *S. luteus* isolates upon 48h exposure to different concentrations of Zn (20, 200, 1000 μM). (a) ZRC1, (b) ZnT1, (c) ZnT2, (d) MT, (e) GSS, (f) Mn-SOD, (g) HSP70 and (h) GST expression. Higher Zn concentrations correspond to darker bars. Data are the average of 5 biological replicates +/− S.E.; significant differences (p<0.05) compared to the control are indicated by *.

Individual isolates show a different metallothionein (MT) expression level (fig. 2d; missing in 2 isolates; highly expressed in one) and the expression of the gene is responsive to Zn (2.2x upregulation, 1000 μM Zn) in only one isolate. Four other genes, involved in general stress response and repair were studied. As for the *MT*, variation in expression level is considerable among isolates but independent of Zn tolerance phenotype (fig. 2e,f,g, h). For all genes, responsiveness to Zn was shown in some of the Zn-sensitive isolates. A decrease of GST expression upon exposure to 1000 μM in Lm8, is the only significant effect of Zn on gene expression which could be detected in a Zn-tolerant isolate (fig. 2h)

### Naturally occurring Zn tolerance phenotypes

Following the initial screening of *S. luteus* isolates with contrasting Zn tolerance phenotypes, a second set of experiments using 30 randomly selected isolates was performed. Dose-response experiments revealed that Zn tolerance in *S. luteus* is a continuous trait (fig. 3). EC50 values ranged from 0.94 mM Zn for the most sensitive isolate P1 to 17.81 mM Zn for the most tolerant isolate OF3. Three isolates collected on non-polluted sites show an intermediate to high Zn tolerance level (MG4, EC50 = 4.60 mM Zn; HH5, EC50 = 5.03 mM Zn; and EW2, EC50 = 16.58 mM Zn). Isolates collected on polluted sites all exhibit an intermediate to high Zn tolerance level.

**Figure 3.**
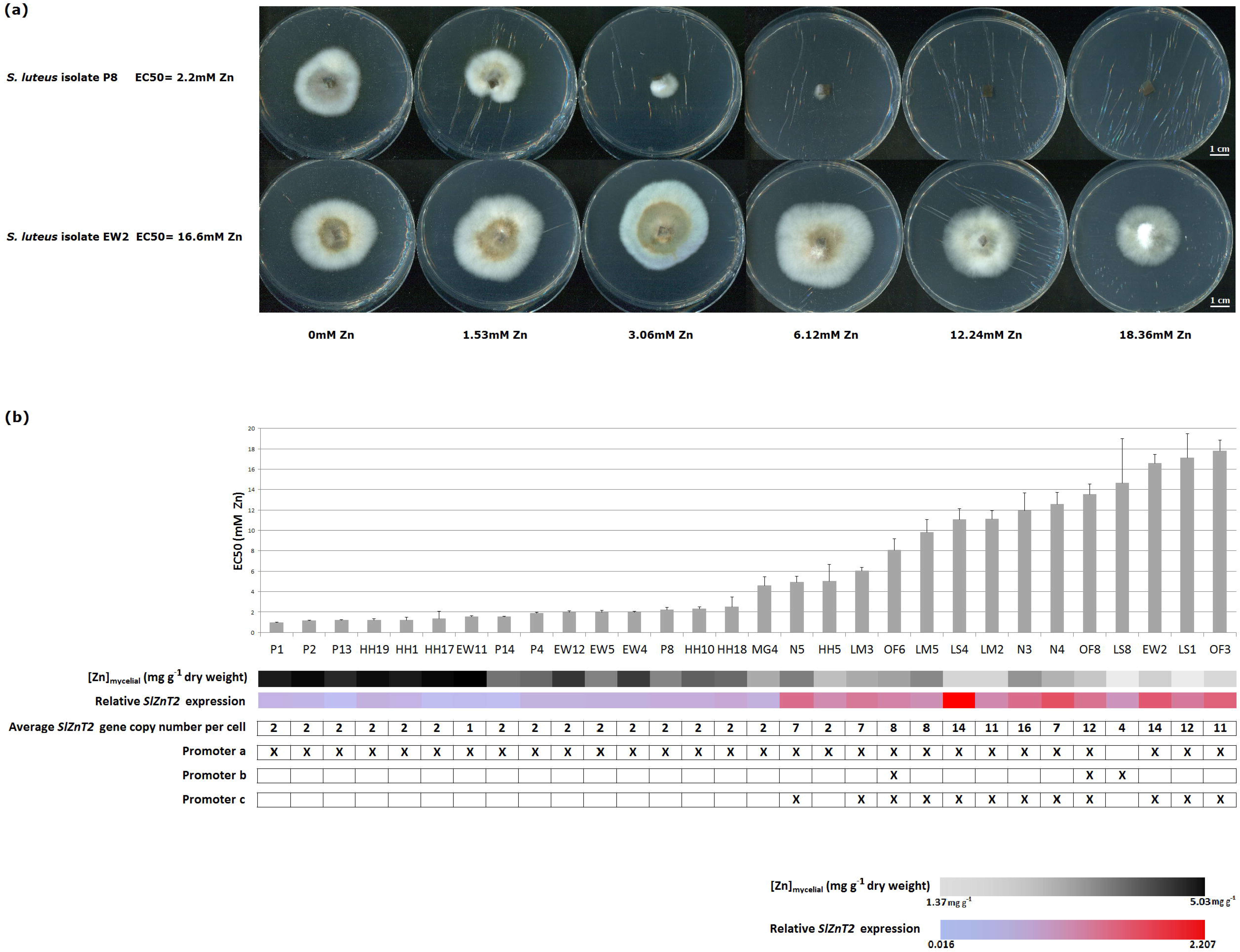
Zn tolerance phenotypes and associated genotypes for different *S. luteus* isolates. (a) Fourteen-day-old *S. luteus* cultures on cellophane-covered petri dishes containing standard modified Fries medium supplemented with 0mM, 1.53mM, 3.06mM, 6.12mM, 12.24mM, and 18.36mM Zn. Upper row: a Zn sensitive isolate P8 (EC50 = 2.20 mM Zn). Lower row: a Zn tolerant isolate EW2 (EC50 = 16.58). (b) Screening of 30 randomly selected *S. luteus* isolates. Isolates are ranked from most sensitive to most tolerant. EC50 values +/− S.E. are displayed in the graph. Mycelial Zn content is visualized for each individual isolate below the graph using a gradient from black to grey, representing the range in Zn content. Relative *SlZnT2* expression is visualized in the next row using a blue to red gradient. The subsequent row states the average *SlZnT2* gene copy number per cell for each isolate. The last 3 rows illustrate the presence (indicated with “X”) of promotor genotype “a”, “b”, and “c” in each isolate.

Following the dose-response experiments, mycelial Zn content was analysed in samples exposed to a relatively low, sublethal Zn treatment of 1.53 mM Zn to ensure sufficient growth of all isolates (Fig. 3 and S3a). A strong negative linear correlation was observed between the EC50 values and mycelial Zn concentrations when both were log transformed (Pearson’s product moment correlation coefficient r = −0.91, r² = 0.83, p < 0.0001; fig. S3b).

### SlZnT2 transcription and correlation with phenotypes

Gene expression of *SlZnT2* was analysed in the larger study group of 30 isolates. When grown in control conditions, fold changes in expression level of up to 137 x were detected among isolates (P13 = 0.016 and Ls4 = 2.20; fig. 3 and S4a). As observed in the first set of experiments, *SlZnT2* transcripts were almost absent in some of the most Zn sensitive isolates (P13, P14, and HH17) whereas it was abundant in most tolerant isolates (EW2, LS1 and OF3). On average an eight-fold difference in expression level was detected when comparing the 6 most Zn sensitive isolates (EC50 < 1.53mM Zn) to the 6 most Zn tolerant isolates (EC50 > 12.24mM Zn). The positive correlation between EC50 value and *SlZnT2* expression is visualized in Fig. S4b. Both variables were log-transformed and Pearson’s product moment correlation confirmed a significant linear relationship (r = 0.84, r² = 0.72, p < 0.0001).

### SlZnT2 copy number variation among isolates

*SlZnT2* gene copy number varied among different *S. luteus* isolates. Gene copy numbers ranged from 0.5 to 8 (N3) per nucleus, i.e. 1 (EW11) to 16 (N3) per cell since all isolates were dikaryotic (fig. 3 and S5a). Isolates with a single copy of the gene show a low to intermediate Zn tolerance level. In highly Zn tolerant isolates, *SlZnT2* is a multi-copy gene. *SlZnT2* gene expression level correlates positively with copy number (Pearson’s product moment correlation coefficient r = 0.80, r² = 0.64, p < 0.0001; fig. S5b). A significant positive correlation between EC50 values and gene copy number was also obvious (Fig. S5c).

### Variation in SlZnT2 promoter genotype among isolates

Upstream region of the *SlZnT2* gene was amplified for 30 *S. luteus* isolates and resulted in 44 sequences. Sequences were submitted to NCBI Genbank (accession numbers MK395391 - 395434). For all 30 *S. luteus* isolates, at least one upstream sequence was identified. All 44 detected sequences show eukaryotic core regulatory elements, i.e. a TATA-box, CCAAT-box and GC-box and can be considered as promoter sequences (Fig. S6). Sequence alignment and similarity analysis grouped the different promoter sequences in three distinct clusters (Fig. 4). Sequences clustering together were considered as the same promoter genotype. Promoter genotypes were named “Promoter a”, “Promoter b” and “Promoter c” according to the cluster they belong to. The clusters are well-supported both by high bootstrap values (100% for each node; 1000 replicates) and branch lengths. Promoter a-like sequences were detected in 29 of the 30 isolates (absent only in LS8, EC50 = 14.64; Fig. 3). Promoter c was detected in 12 of the more Zn tolerant isolates (EC50 ≥ 6.06mM Zn) and Promoter b in three highly Zn tolerant isolates (OF6, EC50 = 8.05mM Zn; OF8, EC50 = 13.54mM Zn; OF6, EC50 = 14.64mM Zn). All three promoter genotypes are characterized by SNPs (single-nucleotide polymorphisms), INDELs (insertions or deletions) and multiple repetitive elements (Fig. S6). Sequence variability was highest in Promoter b. This is reflected in the average branch length within each cluster (Promoter a: 0.008, Promoter b: 0.052, and Promoter c: 0.003).

**Figure 4.**
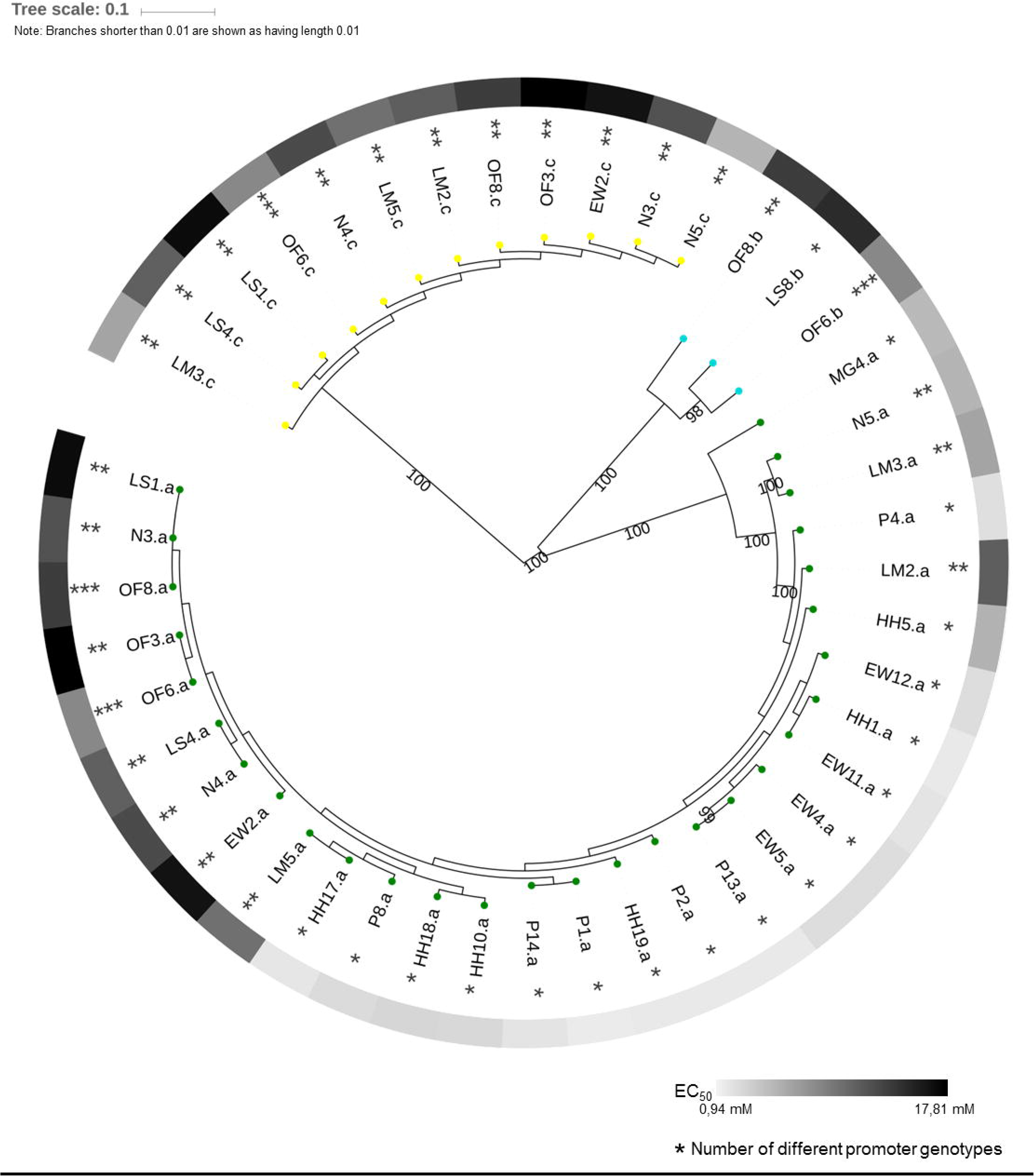
Hierarchical clustering of isolated *SlZnT2* promoter sequences. The 44 identified sequences cluster in three distinct groups: ‘promotor a’ (green), ‘promotor b’ (blue), and ‘promotor c’ (yellow). Zn tolerance phenotype of the *S. luteus* isolate from which they were amplified is indicated by a grey to black gradient. Each sequence is marked with one (*), two (**), or three asterisks (***) to indicate the total number of different promoter sequences identified in that isolate. The UPGMA tree construction method and the Jukes-Cantor distance correction model were used for the construction of the tree. Bootstrap values were estimated from 1000 replicates.

## Discussion

### Common mechanisms of homeostasis and detoxification are differentially regulated among isolates

A limited amount of *S. luteus* isolates with contrasting Zn tolerance phenotypes were studied on different levels after Zn exposure to identify the strategies involved in detoxification of this metal. The activity of anti-oxidative enzymes and lipid peroxidation levels were measured after 48h of exposure to sub-lethal Zn concentrations to reflect early events of cellular stress and damage. There were no indications of accumulation of reactive oxygen species (increased CAT or SOD activity) nor indications of oxidative membrane damage (increased TBARS) in any of the isolates (fig.1). SOD activity decreased in response to Zn in all Zn sensitive isolates. A decrease in SOD activity in response to metal treatment is not common. Mostly, metals provoke an increase in SOD activity to counteract induced oxidative stress (Vallino *et al.*, 2009). Nevertheless, an inhibition of specific SOD isozymes in response to Cd was found in *Rhizopogon luteolus* (Miszalski *et al.*, 1996). Excess Zn might have the same kind of effect in *S. luteus* by causing a (relative) deficiency of co-factor (Mn/Fe) or by competition between Zn and Mn/Fe at co-factor binding sites. Some Zn tolerant *S. luteus* isolates accumulate Fe upon prolonged Zn exposure (Colpaert et al., 2004), likely in an attempt to restore cellular Zn/Fe ratio.

Gene expression of eight genes with putative roles in Zn homeostasis and metal detoxification pathways was profiled. For most of the profiled genes, variation in expression level among isolates with the same Zn tolerance phenotype is as high as variation among isolates with contrasting Zn tolerance phenotypes. Common mechanisms of metal detoxification (sequestration in compartments or by sulphur containing compounds) and repair (heat shock proteins), described for many ectomycorrhizal fungi (Schützendubel & Polle, 2002; Bolchi *et al.*, 2011; Ruytinx et al., 2016) seem to be differentially regulated and deployed among *S. luteus* isolates upon exposure to high external Zn concentrations. A high variation in transcriptomic responses among isolates might contribute to the potential of ectomycorrhizal species to adapt to changing environments. Also *Pisolithus albus*, an ectomycorrhizal fungus of which different Ni tolerance phenotypes exist, shows a high variability in transcriptomic responses towards external Ni concentrations (Majorel et al., 2012).

A clear response of gene expression on external Zn concentration was observed in all *S. luteus* isolates for the Zn importer SlZRT1. As reported by Coninx et al. (2017), transcripts of this gene are present in low amounts when environmental Zn is high. Surprisingly, reaching a similar *SlZRT1* transcript level requires a shift in external concentration when comparing *S. luteus* isolates of contrasting Zn tolerance phenotypes. This could indicate a differential regulation of homeostatic mechanisms among *S. luteus* isolates or more likely that transcript levels of the Zn importer are dependent on intracellular Zn status. The latter suggests the Zn sensor regulating long term homeostatic mechanisms of *S. luteus* in response to environmental changes to be located internally rather than being integrated at the cellular membrane. Zn sensing and regulation of mechanisms involved in Zn homeostasis are not yet clear in filamentous fungi. In yeasts however, environmental Zn sensing leading to adjustment of main homeostatic mechanisms is hypothesized to occur intracellular by direct interaction of Zn and transcriptional activators (Choi & Bird, 2014). As for *SlZRT1*, we detected a difference in transcript level among both Zn tolerance groups of *S. luteus* for *SlZnT2*, a putative Zn transporter of the CDF family. Transcript level is high in Zn tolerant isolates and near to background in Zn sensitive isolates. For nine out of the ten isolates tested, transcript level was not responsive to external Zn concentrations. Therefore, differences in *SlZnT2* transcript level are expected to be due to differences in genetic background among isolates with a different Zn tolerance phenotype rather than being the result of environmental factors. As a result, *SlZnT2* gene expression is a well suited candidate parameter to test in natural populations in an attempt to unravel the genetics of adaptive Zn tolerance phenotypes.

### Zn tolerance is a continuous trait and correlates with SlZnT2 transcript level

Adaptation of *S. luteus* to high environmental Zn concentrations is expected to have occurred rapidly. Zn smelters in Nort-Eastern Belgium polluted the surrounding environment from 1870-1970. *S. luteus* populations were sampled 25-30 years after ending the release of Zn ores, fumes and dust in the region (Colpaert et al., 2004). Nevertheless, despite the recent environmental shift on evolutionary time scale and steep gradient in soil Zn availability, we showed Zn tolerance in *S. luteus* to be a quantitative trait. Phenotypes covering the continuum from extreme Zn sensitive (EC50 = 0.94 mM Zn) to hypertolerant (EC50 = 17.81 mM Zn) were found. Intermediate to high zinc tolerant phenotypes of *S. luteus* were collected at low frequency (2/15 isolates and 1/15 isolates respectively) from non-polluted sample sites. Whether these reflect standing variation within the original population or are due to reciprocal gene flow needs an overall analysis of demographic history. Ongoing gene flow among sample sites and a lack of population structure due to metal availability in soils were demonstrated previously (Muller et al., 2007b).

Quantitative traits are typically controlled by many genetic loci (QTLs or quantitative trait loci) having a cumulative effect and final phenotypes are often the result of genotype x environment interactions (e.g. Frachon et al., 2018; Ferrero-Serrano & Assmann, 2019; Gould et al., 2018). Due to the fact that many genetic loci are contributing to the final tolerance phenotype, the physiological or molecular mechanisms underlying the tolerance trait might be different among individuals. This is the case for Zn and Cd tolerance in *Arabidopsis halleri*. Though Zn/Cd tolerance is a constitutive trait in this plant species, detoxification and accumulation strategy vary among populations (Corso et al., 2018; Schvartzman et al., 2018). We showed *S. luteus* Zn tolerance phenotype to be negatively correlated with intracellular Zn accumulation. No outlier isolates were detected suggesting all individuals to use a similar detoxification strategy based on Zn exclusion as previously demonstrated by Colpaert et al. (2005). The correlation between *SlZnT2* transcript level and Zn tolerance phenotype that we demonstrated in 30 randomly selected *S. luteus* isolates might indicate a role for SlZnT2 in Zn export. Constitutive differences in expression of genes involved in metal detoxification were found to underlie metal tolerance traits in plants and animals (Hanikenne et al., 2008; Roelofs et al., 2009). An elaboration of transformation or genome editing protocols for use in *Suillus* species is needed to confirm the function of SlZnT2 in Zn tolerance.

### Differences in gene expression are due to copy number variation and differences in cis-regulation

Gene regulation depends on multiple genetic and environmental factors. Gene copy number variation (CNV) can result in quantitative variation in gene product and contributes to metal tolerance in hyperaccumulating plants (Hanikenne et al., 2008; Craciun et al., 2012). *SlZnT2* is the result of a gene duplication in the common ancestor of the Suilloid clade and became functionally differentiated from its paralogue *SlZnT1* (Ruytinx et al., 2017). We demonstrated a further multiplication of *SlZnT2* upon speciation in *S. luteus*. Gene duplication events in fungi are common but gene copies, if retained, rarely diverge in biochemical function. Rather they diverge with respect to regulatory control and result in a modified transcript profile (Wapinski et al., 2007). *SlZnT2* gene copy number ranges from 1 - 16 among *S. luteus* isolates and was found to explain 72% of the observed variation in *SlZnT2* transcript accumulation. Whether these copies are tandemly arranged or spread all over the genome remains elusive as well as the mechanisms leading to such a high CNV. Yet, it is clear from Cu tolerance in yeasts that environmental stimulation of CNV allows rapid adaptation (Hull et al., 2017). Given the absence of bottlenecks, ongoing gene flow (Muller et al., 2007) and high CNV among individuals collected at the same sampling site, this might also have occurred and still be ongoing in *S. luteus* isolates thriving on heavy metal contaminated sites in North-East Belgium.

Additional to CNV, cis-regulatory variants are a critical component affecting gene-expression and phenotypic variation (Stern & Orgogozo, 2008). Differences in promoter genotype were detected among different *SlZnT2* gene copies and variation in cis-regulation among copies is expected. Experimental evolution in *Escherichia coli* showed random sequences to evolve rapidly in *de novo* promoters and small sequence differences were shown to result in major differences in expression level (Yona et al., 2018). Three major promoter genotypes could be identified for *SlZnT2* in *S. luteus*. Since all (except one) isolates possess promoter genotype “a” we assume this to be ancestral and the others (“b” and “c”) to be linked to additional acquired SlZnT2 copy’s. Contribution of the individual genotypes to overall expression level is unclear but assumed to be unequal. Promoter genotype “a” is assumed to result in a low basal expression level and the other promoter genotypes to result in a higher transcript level. However, promoter genotype x CNV interactions impede the deduction of effect sizes in the current data-set.

### Conclusion

We showed the cumulative effect of gene CNV and differentiation in promoter genotype to act on the expression of a common gene and to support rapid environmental adaptation. The high variation in gene copy number among individuals sampled at the same geographical site along with a lack of bottlenecks and population structure suggest adaptation through stimulated copy number variation rather than being the result of standing variation. However, to support this hypothesis, an in depth analysis of the considered genetic regions is required among others, to fully understand the genomic features allowing multiple gene copy’s to emerge and accumulate.

## Experimental procedures

### Fungal material and growth conditions

A limited number (10) of isolates of *Suillus luteus* (L.:Fr.) showing extreme Zn tolerance phenotypes, i.e. hypertolerant or hypersensitive, were used for initial experiments aiming at characterizing the physiology of Zn tolerance and identifying candidate genes underlying the Zn tolerance trait. These isolates were collected and categorized according to published EC50-values (Colpaert *et al.*, 2004; Muller *et al.*, 2004). Subsequently 30 *S. luteus* isolates, randomly selected from UHasselt fungal culture collection were used in an experiment aiming at associating Zn tolerance phenotype with a particular genetic factor. An overview of the isolates and their geographical collection sites is provided in TableS1 and Figure S1. All isolates are assumed to be dikaryotes unless stated otherwise.

Isolates were individually cultured on cellophane-covered solid modified Fries medium (28 mM glucose, 5.4 mM ammonium tartrate, 1.5 mM KH_2_PO_4_, 0.4 mM MgSO_4_·7H_2_O, 5 μM CuSO_4_·5H_2_O, 20 μM ZnSO_4_·7H_2_O, 0.1 μM biotin, 0.5 μM pyridoxine, 0.3 μM riboflavin, 0.8 μM nicotinamide, 0.7 μM p-aminobenzoic acid, 0.3 μM thiamine, 0.2 μM Ca-pantothenate and 0.8% agar; pH-adjusted to 4.8). Plates were incubated at 23°C in the dark. One-week old mycelia were used to prepare liquid cultures for the physiology experiment. Two-week old mycelia were subcultured for tolerance assays or harvested for RNA and DNA extraction in the association experiment. Liquid cultures were prepared, incubated for one week and exposed for 48h to 20, 200 or 1000 μM Zn according to Ruytinx et al., 2017. Mycelia were weighed after treatment. Sample aliquots (200 mg and 500 mg) were flash frozen and stored at −70°C for lipid peroxidation, enzyme and gene expression assays.

### Enzyme activities and lipid peroxidation

Frozen mycelia (500 mg) of six *S. luteus* isolates were homogenized in 2 ml ice-cold 0.1 M Tris-HCl buffer (pH 7.8) containing 1 mM EDTA, 1 mM dithiotreitol and 4% insoluble polyvinylpyrrolidone. The homogenate was squeezed through a nylon mesh and centrifuged for 10 min at 19000 g and 4°C. The enzyme activities were measured spectrophotometrically in the supernatant at 25°C. Analysis of superoxide dismutase (SOD, EC 1.15.1.1) activity was based on the inhibition of cytochrome c at 550 nm (McCord & Fridovich, 1969). Catalase (CAT, EC 1.11.1.6) activity was measured at 240 nm as described by Bergmeyer (1974).

Thiobarbituric acid reactive substances (TBARS) were measured spectrophotometrically to estimate the amount of lipid peroxidation and oxidative damage. Frozen mycelia (500 mg) were homogenized in 1 ml 0.1% TCA and 4 ml 0.5% TBA was added to the extract. This mixture was heated to 95°C for 30 min and subsequently centrifuged for 10 min at 19 000 g. Absorbance of the supernatant was measured at 532 nm. The obtained values were corrected for unspecific absorbance by subtracting absorbance reads at 600 nm.

One way ANOVA followed by Dunnett post-hoc test was used to identify significant differences (p < 0.05) as a consequence of metal treatment in single *S. luteus* isolates. Transformations were applied when necessary to approximate normality. All statistical analyses were performed using the SigmaStat 3.2 software package.

### Gene expression analysis of selected candidate genes

Frozen spherical mycelia (200 mg) from liquid cultures were thoroughly ground in liquid nitrogen using a mortar and pestle. Total RNA was extracted from the ground tissue of 10 different *S. luteus* isolates using the RNeasy Plant mini kit (Qiagen). Genomic DNA was eliminated using the TURBO DNA-free kit (Applied Biosystems) and one μg of total RNA was used in a High-Capacity cDNA Reverse Transcription reaction (Applied Biosystems), according to the manufacturer’s instructions. The cDNA was diluted 10x in a tenfold dilution of TE buffer (1mM Tris-HCl, 0.1mM EDTA, pH 8.0) and stored at −20°C.

Expression levels of eight genes (*MnSOD, HSP70, ZRT1, ZnT1, ZnT2, GST, GSS, MTa*) and three reference genes (*GR975621, AM085168, AM085296*) were measured. Primer sequences (Table S2) were taken from Coninx *et al.*, (2017), Muller *et al.* (2007), Nguyen *et al.* (2017), Ruytinx *et al.* (2011, 2016, 2017), or newly designed using the Primer3 program (Rozen & Skaletsky, 2000). Real-time PCR was performed in an optical 96-well plate with an ABI PRISM 7500 sequence detection system (Applied Biosystems) and fast cycling conditions (20s at 95°C, 40 cycles of 3s at 95°C and 30s at 60°C) according to Ruytinx *et al.*, 2017. The number of reference genes and their stability was approved and expression levels were calculated according to Ruytinx *et al.* (2016). Significant differences (p<0.05) in transcript level due to metal treatment in single isolates were evaluated using one-way ANOVA and Dunnett post-hoc tests using the SigmaStat 3.2 software package. Transformations were applied when necessary to approximate normality.

*SlZnT2* transcription for mycelia grown on cellophane-covered plates in the association experiment was determined with minor adjustments to the protocol according to Coninx *et al.* (2017). Reference genes *TUB1, ACT1* and *GR97562* were used for normalization.

### Zn tolerance and accumulation assays

Zn tolerance was evaluated via dose-response experiments. Inocula (0.5 cm² plugs) were transferred to cellophane-covered petri dishes of modified Fries medium enriched with ZnSO_4_·7H_2_O (Colpaert et al., 2004). Zn was added to the growth medium in the following concentrations: 0, 1.5, 3.1, 6.2, 12.3 and 18.5 mM Zn. Metal exposures were performed in five replicates. After two weeks, mycelia were harvested and lyophilized. Samples were weighed to obtain their dry weights and based on these data half maximal effective concentration (EC50) values were calculated for each *S. luteus* isolate. EC50 values were determined via non-linear regression with a four parameter log-logistic model (Ritz et al., 2015) in “R” version 3.5.1 (R Core Team, 2018). EC50 values are represented as the zinc concentrations that inhibit growth by 50% compared to the control condition.

Lyophilized mycelia exposed to 1.5 mM Zn were acid digested (HNO_3_/HCl) (Colpaert et al., 2005) and mycelial Zn concentrations were analysed by inductively coupled plasma optical emission spectrometry (ICP-OES).

### DNA extraction and ZnT2 gene copy number

Fungal mycelia were powdered in liquid nitrogen using mortar and pestle. DNA was extracted using a CTAB/phenol/chloroform protocol (Methods S1; modified from Liao *et al.*, 2014). Samples were processed with the GeneJET PCR Purification Kit (Thermo Fisher Scientific, Waltham, MA, USA) according to manufacturer’s instructions. DNA concentration was determined on a Qubit 2.0 fluorometer (Life Technologies, Carlsbad, CA, USA) with the Qubit dsDNA HS Assay Kit (Invitrogen, Carlsbad, CA, USA) following the manufacturer’s protocol.

*SlZnT2* gene copy number was estimated via qPCR (D’Haene *et al.*, 2010). Reaction mix and cycling condition were identical to those used for gene expression analysis (described above) with the exception of genomic DNA (1.25 ng) serving as PCR template. For each isolate, reactions were run in triplicate using three independent gDNA samples. The *ACT1* gene was used as a calibrator, since all isolates were shown to have an equal (single) copy number of this gene (Fig. S2). *SlZnT2* copy numbers were obtained by calculating the ratio of 2^−ΔCt^ of *SlZnT2* and *ACT1*. Two sets of primer pairs, spanning intron-exon boundaries, were designed for *ACT1* and *SlZnT2* using Primer3web version 4.1.0 (Rozen and Skaletsky, 2000). For each gene measurement the average value of the two primer pairs (Table S2) was used to minimalize effects of differences in primer pair efficiency among isolates.

### Genome walking and ZnT2 promoter analysis

DNA of six *S. luteus* isolates was used in a genome walking procedure using the Universal GenomeWalker™ kit (Clontech, Mountain View, California, US) according to the manufacturer’s instructions. Gene-specific primers GSP1 (5’-TGA AGA TGT GGC TCA CCG ACG ACG AGT-3’) and GSP2 (5’-GTG ATG CGA GCT GAA CGA GAT AAT GCC AT-3’) were developed for the primary and nested PCR, respectively. Primers were designed with Primer3web version 4.1.0 software (Rozen and Skaletsky, 2000). PCR amplicons were visualized via gel electrophoresis with GelRed® Nucleic Acid Gel Stain (Biotium, Fremont, California, US) and processed with the QIAquick Gel Extraction Kit (Qiagen, Hilden, Germany). PCR products were cloned into the pCR™4-TOPO® TA vector (Invitrogen, Carlsbad, California, US) and sent for sequencing (Macrogen, Amsterdam, The Netherlands). Promoter sequences were assembled and aligned with CLC Main Workbench 8 software (Qiagen, Hilden, Germany).

Primers for the PCR amplification of these promoter sequences in 30 *S. luteus* isolates were designed and are listed in Table S2. Each reverse primer is located on the *SlZnT2* gene. The Platinum Taq DNA polymerase high fidelity kit (Invitrogen, Carlsbad, California, US) was used according to the manufacturer’s instructions. PCR cycling conditions are specified in Methods S2. PCR products were visualised via gel electrophoresis, gel extracted, and sequenced (Macrogen, Amsterdam, The Netherlands). Sequences were analysed with GPMiner (Gene Promoter Miner) [accessed: 19 August 2018] (Lee et al., 2012) for the presence of core regulatory features of promoter sequences (TATA-box, CCAAT-box, and GC-box). Afterwards, the entire set of promoter sequences was aligned and a phylogenetic tree was constructed with CLC Main Workbench 8. The phylogenetic tree was constructed via the UPGMA (Unweighted Pair Group Method with Arithmetic Mean) method and the Jukes-Cantor distance correction model and exported to iTOL v4.3 (Letunic & Bork, 2016) for further annotation.

### Association analysis

Data were statistically analysed with “R” version 3.5.1 (R Core Team, 2018). Data were log-transformed and data normality was verified with the Shapiro-Wilks test. Heteroscedasticity was evaluated with the Breusch-Pagan test. Correlation analyses were performed with Pearson product-moment correlation. Since multiple comparisons were made, p-values were adjusted with the Bonferroni correction.

## Acknowledgements

We thank Ann Wijgaerts for assistance in DNA extractions, dr. Bianca Cox for statistical advice and Carine Put for maintaining the *Suillus* culture collection. This work was financially supported by the Research Foundation – Flanders (FWO-project G.0925.10N and G079213). A Flanders Innovation & Entrepreneurships PhD fellowship (IWT project 141461) was assigned to Laura Coninx. The authors declare no conflicts of interest.

---

**Figure S1. Geographical map indicating the *S. luteus* sampling sites in the Limburg province in Belgium**. *S. luteus* sampling sites are indicated with a blue “x”. The full names of the sampling sites are given in the legend. Zn smelters in the region are indicated by a brown factory symbol.

**Figure S2. Average gene copy number of *ACT1* +/− SE of three biological replicates.** The sequenced monokaryotic isolate UH-LM8-n1 was included as a control (indicated in green), since it is known to have one gene copy of *ACT1*. The *ACT1* copy numbers are expressed as copy number per nucleus, since all isolates except from the control result from sporocarps and are assumed to be dikaryotic.

**Figure S3. Mycelial Zn concentrations and correlation with Zn tolerance.** (a) Average mycelial Zn concentrations +/− SE of five biological replicates. *S. luteus* isolates were exposed to 1.53mM Zn for 14 days and are displayed from most sensitive (left) to most tolerant to Zn (right). (b) Significant negative linear relationship between log-transformed EC50-values and log-transformed mycelial Zn concentrations of *S. luteus* isolates. (Pearson’s product-moment correlation r = −0.91, r² = 0.83, p < 0.001).

**Figure S4. Relative *SlZnT2* gene expression and correlation with Zn tolerance.** (a) Relative *SlZnT2* gene expression +/− SE of three biological replicates. Isolates are arranged from most sensitive (left) to most tolerant to Zn (right). (b) 4 Significant positive linear relationship between log-transformed EC50-values and log-transformed *SlZnT2* transcript levels in *Suillus luteus* isolates. (Pearson’s product-moment correlation r = 0.84, r² = 0.72, p < 0.001).

**Figure S5. *SlZnT2* gene copy numbers and correlation** with ***SlZnT2* expression and Zn tolerance.** (a) Average *SlZnT2* gene copy number per nucleus +/− SE of three biological replicates. As all isolates included in the study are assumed to be dikaryotic, values need to be doubled to represent the average *SlZnT2* copy number per cell. Isolates are displayed from most sensitive (left) to most tolerant to Zn (right). (b) Significant positive linear relationship between log-transformed *SlZnT2* copy numbers and log-transformed *SlZnT2* transcript levels in *Suillus luteus* isolates. (Pearson’s product-moment correlation r = 0.80, r² = 0.64, p < 0.001). (c) Correlation analysis of the log-transformed EC50 values and *SlZnT2* gene copy numbers (Pearson’s product-moment correlation r = 0.88, r² = 0.77, p < 0.0001).

**Figure S6. Consensus sequences of the promotor genotypes with core regulatory elements.** (a) promotor a, (b) promotor b, and (c) promotor c. TATA boxes (green), CAAT boxes (blue), and GC boxes (red) are indicated on the sequences.

## Supplementary Table legends

**Table S1** Primer sequences of primers used for gene expression measurements, copy number assessment, and *SlZnT2* promotor amplification.

**Table S2** Genes of interest description and JGI Protein ID numbers.

**Table S3** Overview of all the *S. luteus* isolates included in this study together with their geographical collection sites. All collection sites were located in the province Limburg (Belgium). Characteristics of the isolates and collection sites were described previously (Colpaert et al., 2004; Muller et al., 2004).

## References

Adriaensen K, Vrålstad T, Noben JP, Vangronsveld J, Colpaert JV. 2005. Copper-adapted Suillus luteus, a symbiotic solution for pines colonizing Cu mine spoils. Applied and Environmental Microbiology 71: 7279–7284.

Anderson JT, Willis JH, Mitchell-Olds T. 2011. Evolutionary genetics of plant adaptation. Trends Genet. 27(7):258–266.

Bergmeyer HU, Gawenn K, Grassl M. 1974. Enzymes as Biochemical Reagents. In: Bergmeyer HU, eds. Methods in Enzymatic Analysis. New York: Academic Press, 425–522.

Bolchi A, Ruotolo R, Marchini G, Vurro E, Sanita di Toppi L, Kohler A, Tisserant E, Martin F, Ottonello S. 2011. Genome-wide inventory of metal homeostasis-related gene products including a functional phytochelatin synthase in the hypogeous mycorrhizal fungus *Tuber melanosporum*. Fungal Genetics and Biology 48: 573–584.

Bradford MA, McCulley RL, Crowther TW, Oldfield EE, Wood SA, Fierer N. 2019. Cross-biome patterns in soil microbial respiration predictable from evolutionary theory on thermal adaptation. Nature Ecology & Evolution. doi:10.1038/s41559-018-0771-4.

Branco S, Gladieux P, Ellison CE, Kuo A, Labutti K, Lipzen A, Grigoriev IV, Liao H, Vilgalys R, Peay KG, Taylor JW, Bruns TD. 2015. Genetic isolation between two recently diverged populations of a symbiotic fungus. Molecular Ecology. 24(11): 2747–2758.

Branco S, Bi S, Liao H, Gladieux P, Badouin H, Ellison CE, Nguyen NH, Vilgalys R, Peay KG, Taylor JW, Bruns TD. 2017. Continental‐level population differentiation and environmental adaptation in the mushroom Suillus brevipes. Molecular Ecology 26(7): 2063–2076.

Castaño C, Lindahl BD, Alday JG, Hagenbo A, Martinez de Aragón J, Parladé J, Pera J, Bonet JA.2018. Soil microclimate changes affect soil fungal communities in a Mediterranean pine forest. New Phytol. 220(4):1211–1221.

Choi S, Bird AJ. 2014. Zinc’ing sensibly: controlling zinc homeostasis at the transcriptional level. Metallomics. 6(7): 1198–1215.

Colpaert JV, Muller LAH, Lambaerts M, Adriaensen K, Vangronsveld J. 2004. Evolutionary adaptation to Zn toxicity in populations of Suilloid fungi. New Phytologist 162(2): 549–559.

Colpaert JV, Adriaensen K, Muller LAH, Lambaerts M, Faes C, Carleer R, Vangronsvel J. 2005. Element profiles and growth in Zn-sensitieve and Zn-resistant Suilloid fungi. Mycorrhiza 15: 628–634.

Colpaert JV, Wevers J, Krznaric E, Adriaensen K. 2011. How metal-tolerant ecotypes of ectomycorrhizal fungi protect plants from heavy metal pollution. Annals of Forest Science 68(1): 17–24.

Coninx L, Thoonen A, Slenders E, Morin E, Arnauts N, Op De Beeck M, Kohler A, Ruytinx J, Colpaert JV. 2017. The SlZRT1 Gene Encodes a Plasma Membrane-Located ZIP (Zrt-, Irt-Like Protein) Transporter in the Ectomycorrhizal Fungus Suillus luteus. Front Microbiol 8:2320.

Corso M, Schvartzman MS, Guzzo F, Souard F, Malkowski E, Hanikenne M, Verbruggen N.2018. Contrasting cadmium resistance strategies in two metallicolous populations of Arabidopsis halleri. New Phytologist. 218(1): 283–297.

Craciun AR, Meyer C-L, Chen J, Roosens N, De Groodt R, Hilson P, Verbruggen N.2012. Variation in *HMA4* gene copy number and expression among *Noccaea caerulescens* populations presenting different levels of Cd tolerance and accumulation. Journal of Experimental Botany 63: 4179–4189.

Daborn PJ, Yen JL, Bogwitz MR, Le Goff G, Feil E, Jeffers S, Tijet N, Perry T, Heckel D, Batterham P, Feyereisen R, Wilson TG, ffrench-Constant RH. 2002. A single p450 allele associated with insecticide resistance in Drosophila. Sience. 297(5590): 2253–2256.

D’haene B, Vandesompele J, Hellemans J.2010. Accurate and objective copy number profiling using real-time quantitative PCR. Methods. 50: 262–270.

Dickie IA, Bufford JL, Cobb RC, Desprez-Loustau ML, Grelet G, Hulme PE, Klironomos J, Makiola A, Nuñez MA, Pringle A, Thrall PH, Tourtellot SG, Waller L, Williams NM. 2017. The emerging science of linked plant-fungal invasions. New Phytol. 215(4): 1314–1332.

Fernandez AL, Sheaffer CC, Wyse DL, Staley C, Gould TJ, Sadowsky MJ. 2016. Associations between soil bacterial community structure and nutrient cycling functions in long-term organic farm soils following cover crop and organic fertilizer amendment. Science of the Total Environment. 566-567: 949–959.

Ferrero-Serrano A, Assmann SM. 2019. Phenotypic and genome-wide association with the local environment of Arabidopsis. Nat Ecol Evol. doi: 10.1038/s41559-018-0754-5.

Frachon L, Bartoli C, Carrère S, Bouchez O, Chaubet A, Gautier M, Roby D, Roux F. 2018. A Genomic Map of Climate Adaptation in Arabidopsis thaliana at a Micro-Geographic Scale. Front Plant Sci. 9: 967. doi: 10.3389/fpls.2018.00967.

Franklin J, Serra-Diaz JM, Syphard AD, Regan HM. 2016. Global change and terrestrial plant community dynamics. Proc Natl Acad Sci USA. 113(14): 3725–3734.

Gould BA, Chen Y, Lowry DB. 2018. Gene regulatory divergence between locally adapted ecotypes in their native habitats. Mol Ecol. 27(21): 4174–4188.

Hanikenne M, Talke IN, Haydon MJ, Lanz C, Nolte A, Motte P, Kroymann J, Weigel D, Krämer U. 2008. Evolution of metal hyperaccumulation required cis-regulatory changes and triplication of HMA4. Nature 453: 391–395.

Hull RM, Cruz C, Jack CV, Housely J. 2017. Environmental change drives accelerated adaptation through stimulated copy number variation. PLOS Biology 15(6): e2001333.

Krznaric E, Verbruggen N, Wevers JHL, Carleer R, Vangronsveld J, Colpaert JV. 2009. Cd-tolerant Suillus luteus: a fungal insurance for pines exposed to Cd. Environ Pollut 157: 1581–1588.

Hayward J, Horton TR, Pauchard A, Nunez MA. 2015. A single ectomycorrhizal fungal species can enable a Pinus invasion. Ecology 96: 1438–1444.

Johnson D, Anderson IC, Williams A, Whitlock R, Grime JP. 2010. Plant genotypic diversity does not beget root-fungal species diversity. Plant and Soil. 336(1-2): 107–111.

Jourand P, Ducousso M, Loulergue-Majorel C, Hannibal L, Santoni S, Prin Y, Lebrun M. 2010. Ultramafic soils from New Caledonia structure Pisolithus albus in ecotype. FEMS Microbiology Ecology 72: 238–249.

Jourand P, Hannibal L, Majorel C, Mengant S, Ducousso M, Lebrun M. 2014. Ectomycorrhizal Pisolithus albus inoculation of Acacia spirorbis and Eucalyptus globulus grown in ultramafic topsoil enhances plant growth and mineral nutrition while limits metal uptake. Journal of Plant Physiology. 171(2): 164–172.

Kyaschenko J, Clemmensen KE, Hagenbo A, Karltun E, Lindahl BD. 2017. Shift in fungal communities and associated enzyme activities along an age gradient of managed Pinus sylvestris stands. ISME J. 11(4): 863–874.

Lee TY, Chang WC, Hsu JBK, Chang TH, Shien DM. 2012. GPMiner: an integrated system for mining combinatorial cis-regulatory elements in mammalian gene group. BMC Genomics. 13(1):S3. doi:10.1186/1471-2164-13-S1-S3.

Letunic I, Bork P. 2016. Interactive tree of life (iTOL) v3: an online tool for the display and annotation of phylogenetic and other trees. Nucleic Acids Res. 44(W1): W242–5.

Liao HL, Chen y, Bruns TD, Peay KG, Taylor JW, Branco S, Talbot JM, Vilgalys R. 2014. Metatranscriptomic analysis of ectomycorrhizal roots reveal genes associated with Piloderma-Pinus symbiosis: Improved methodologies for assessing gene expression in situ. Environ Microbiol. 16(12): 3730–3742.

Mack KL, Ballinger MA, Phifer-Rixey M, Nachman MW. 2018. Gene regulation underlies environmental adaptation in house mice. Genome Res. 28(11): 1636–1645.

Majorel C, Hannibal L, Soupe ME, Carriconde F, Ducousso M, Lebrun M, Jourand P. 2012. Tracking nickel-adaptive biomarkers in *Pisolithus albus* from New Caledonia using a transcriptomic approach. Molecular Ecology 21: 2208–2223.

McCord J, Fridovich I. 1969. Superoxide dismutase: an enzymatic function for erythrocuperein (Hemocuprein). The Journal of Biological Chemistry 244: 6049–6055.

Mizalski Z, Botton B, Turnau K. 1996. New SOD isoforms in *Rhizopogon roseolus* (Corda in Sturm) in the presence of cadmium. Acta Physiologiae Plantarum 18: 129–134.

Muller LAH, Lambaerts M, Vangronsveld J, Colpaert JV. 2004. AFLP-based assessment of the effects of environmental heavy metal pollution on the genetic structure of pioneer populations of *Suillus luteus*. New Phytologist 164:297–303.

Muller LAH, Craciun AR, Ruytinx J, Lambaerts M, Verbruggen N, Vangronsveld J, Colpaert JV. 2007. Gene expression profiling of a Zn-tolerant and a Zn-sensitive *Suillus luteus* isolate exposed to increased external zinc concentrations. Mycorrhiza 17: 571–580.

Muller LAH, Vangronsveld J, Colpaert JV. 2007b. Genetic structure of *Suillus luteus* populations in heavy metal polluted and nonpolluted habitats. Molecular Ecology 16: 4728–4737.

Naranjo S, Smith JD, Artieri CG, Zhang M, Zhou Y, Palmer ME, Fraser HB. 2015. Dissecting the Genetic Basis of a Complex cis-Regulatory Adaptation. PLoS Genet. 11(12): e1005751.

Nguyen H, Rineau F, Vangronsveld J, Cuypers A, Colpaert JV, Ruytinx J. 2017. A novel, highly conserved metallothionein family in basidiomycete fungi and characterization of two representative SlMTa and SlMTb genes in the ectomycorrhizal fungus Suillus luteus. Environ. Microbiol. 19, 2577–2587. doi: 10.1111/1462-2920.13729.

Op De Beeck M, Lievens B, Busschaert P, Rineau F, Smits M, Vangronsveld J, Colpaert JV. 2015. Impact of metal pollution on fungal diversity and community structures. Environmental Microbiology. 17(6): 2035–2047.

Phifer-Rixey M, Bi K, Ferris KG, Sheehan MJ, Lin D, Mack KL, Keeble SM, Suzuki TA, Good JM, Nachman MW. 2018. The genomic basis of environmental adaptation in house mice. PLoS Genet. 14(9): e1007672.

Policelli N, Bruns TD, Vilgalys R, Nuñez MA. 2019. Suilloid fungi as global drivers of pine invasions. New Phytol. Accepted Author Manuscript. doi:10.1111/nph.15660.

R Core Team. 2018. R: A language and environment for statistical computing. R Foundation for Statistical Computing, Vienna, Austria. [WWW document] URL https://www.R-project.org/. [accessed 1 August 2018].

Roelofs D, Janssens TKS, Timmermans MJTN, Nota B, Mariën J, Bochdanovits Z, Ylstra B, van Straalen MN. 2009. Adaptive differences in gene expression associated with heavy metal tolerance in the soil arthropod *Orchesella cincta*. Molecular Ecology 18: 3227–3239.

Rozen S, Skaletsky HJ. 2000. Primer 3 on the www for general users and for biologist programmers. In: Krawetz S, Misener S, eds. Bioinformatics Methods and Protocols: Methods in Molecular Biology. Totowa, NJ: Humana Press, 365–386.

Ruytinx J, Craciun AR, Verstraelen K, Vangronsveld J, Colpaert JV, Verbruggen N. 2011. Transcriptome analysis by cDNA-AFLP of *Suillus luteus* Cd-tolerant and Cd-sensitive isolates. Mycorrhiza 21: 145–154.

Ruytinx J, Remans T, Colpaert JV. 2016. Gene expression studies in different genotypes of an ectomycorrhizal fungus require a high number of reliable reference genes. PeerJ Preprints 4:e2125v1 https://doi.org/10.7287/peerj.preprints.2125v1.

Ruytinx J, Coninx L, Nguyen H, Smisdom N, Morin E, Kohler A, Cuypers A, Colpaert J. 2017. Identification, evolution and functional characterization of two Zn CDF-family transporters of the ectomycorrhizal fungus *Suillus luteus*. Environmental Microbiology Reports 9(4): 419–427.

Selby JP, Willis JH. 2018. Major QTL controls adaptation to serpentine soils in Mimulus guttatus. Molecular Ecology 27(24): 5073–5087.

Schvartzman MS, Corso M, Fataftah N, Scheepers M, Nouet C, Bosman B, Carnol M, Motte P, Verbruggen N, Hanikenne M. 2018. Adaptation to high zinc depends on distinct mechanisms in metallicolous populations of Arabidopsis halleri. New Phytologist. 218(1): 269–282.

Stern DL, Orgogozo V. 2008. The loci of evolution: How predictable is genetic evolution? Evolution, 62: 2155–2177. doi:10.1111/j.1558-5646.2008.00450.x

Taylor JW, Branco S, Gao C, Hann-Soden C, Montoya L, Sylvain I, Gladieux P. 2017. Microbiol Spectr. 5(5) doi: 10.1128/microbiolspec.FUNK-0057-2016.

Vallino M, Martino E, Boella F, Murat C, Chiapello M, Perotto S. 2009. Cu,Zn superoxide dismutase and zinc stress in the metal-tolerant ericoid mycorrhizal fungus *Oidiodendron maius* Zn. FEMS Microbiology Letters 293: 48–57.

van’t Hof AE, Campagne P, Rigden DJ, Yung Cj, Lingley J, Quail MA, Hall N, Darby AC, Saccheri IJ. 2016. The industrial melanism mutation in British peppered moths is a transposable element. Nature 534: 102–105.

Wang J, Chen C, Ye Z, Li J, Feng Y, Lu Q. 2018. Relationships Between Fungal and Plant Communities Differ Between Desert and Grassland in a Typical Dryland Region of Northwest China. Front Microbiol 9: 2327. doi: 10.3389/fmicb.2018.02327.

Wapinski I, Pfeffer A, Friedman N, Regev A. 2007. Natural history and evolutionary principles of gene duplication in fungi. Nature 449: 54–64.

Wardle DA, Bardgett RD, Klironomos JN, Setälä H, van der Putten WH, Wall DH. 2004. Ecological linkages between aboveground and belowground biota. Science. 304(5677): 1629–1633.

Yona AH, Alm EJ, Gore J. 2018. Random sequences rapidly evolve into de novo promoters. Nature communications. 9: 1530.

Zhang H, Cai Y, Li X, Christie P, Zhang J, Gai J. 2019. Temperature-mediated phylogenetic assemblage of fungal communities and local adaptation in mycorrhizal symbioses. Environmental Micorbiology Reports doi:10.1111/1758-2229.12729.

