## Supplementary figures and images for "Adaptive zinc tolerance is supported by extensive gene multiplication and differences in cis-regulation of a CDF transporter in an ectomycorrhizal fungus"

### Figure S1

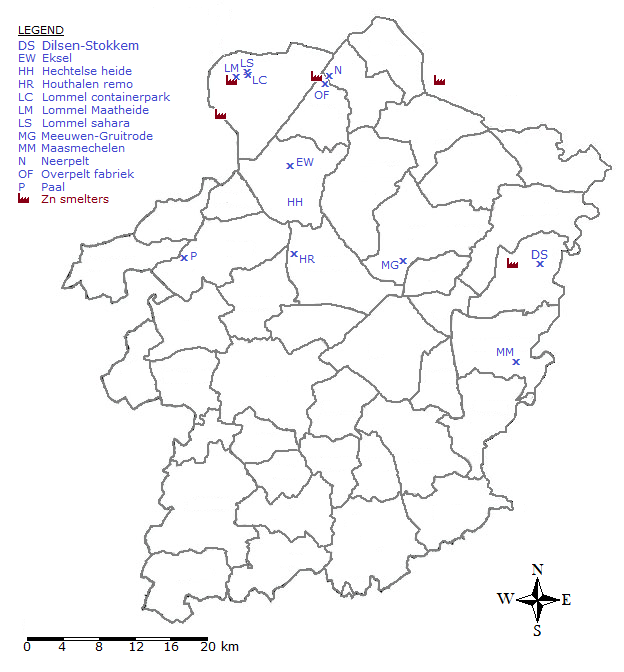

### Figure S2

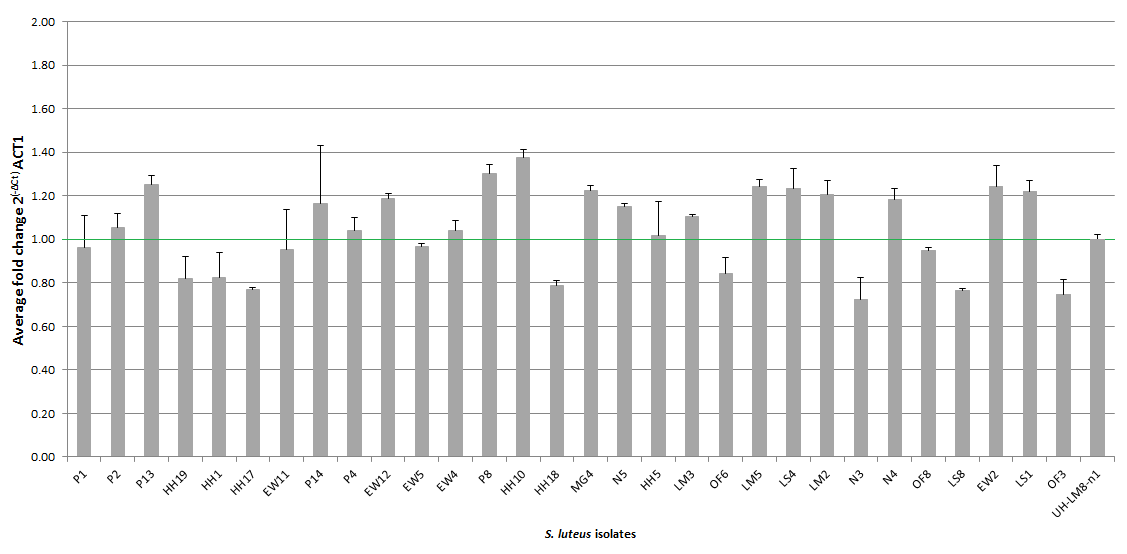

### Figure S3

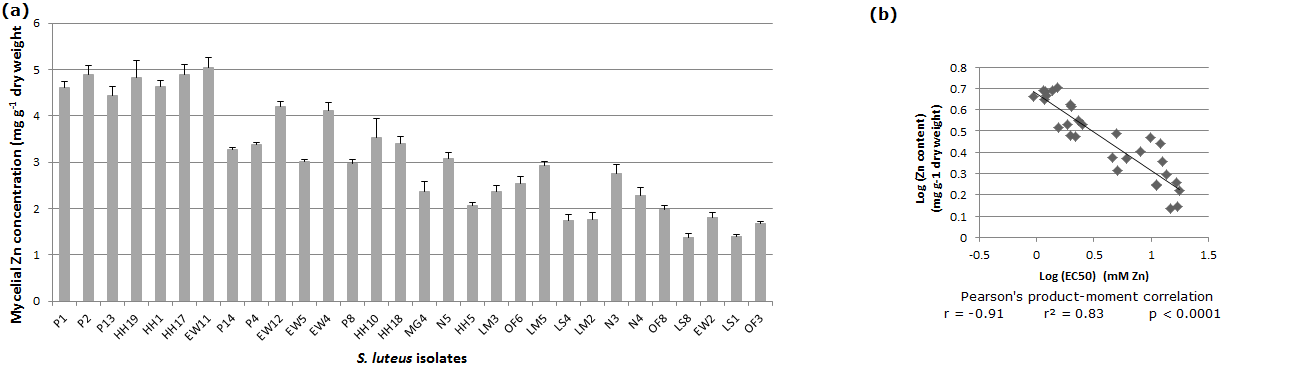

### Figure S4

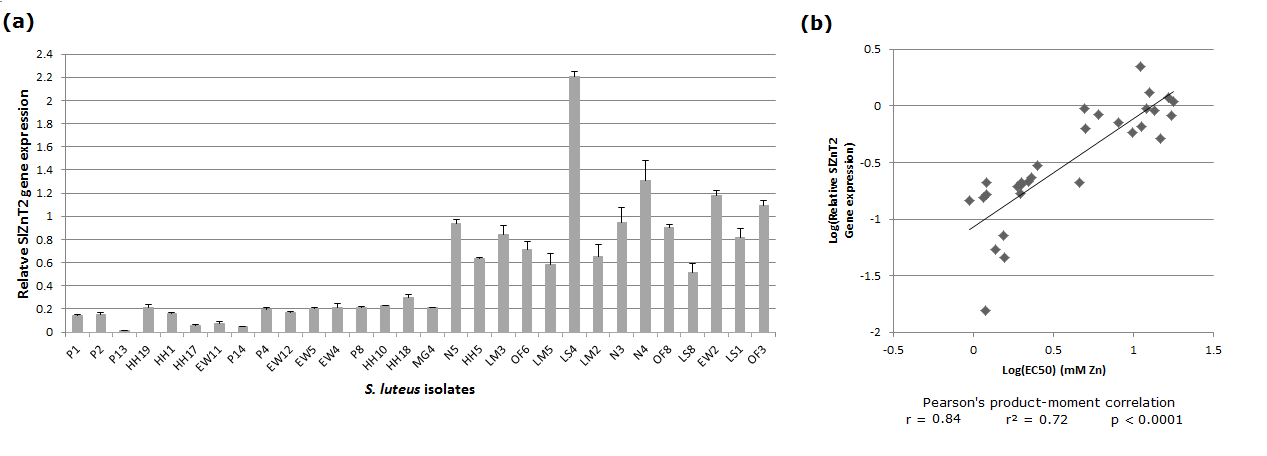

### Figure S5

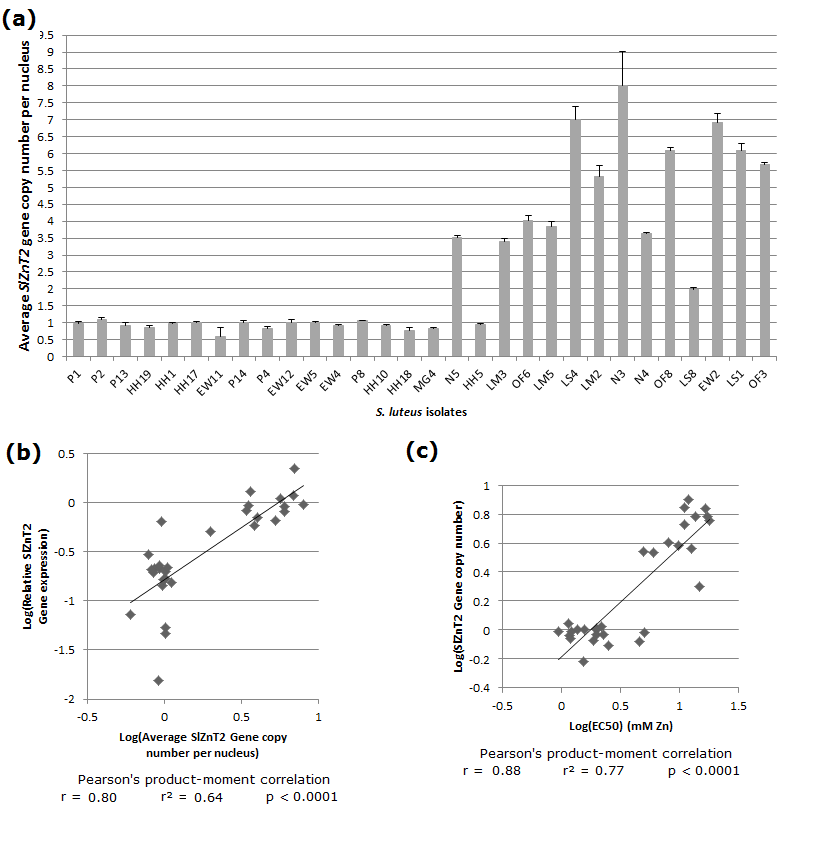

### Figure S6

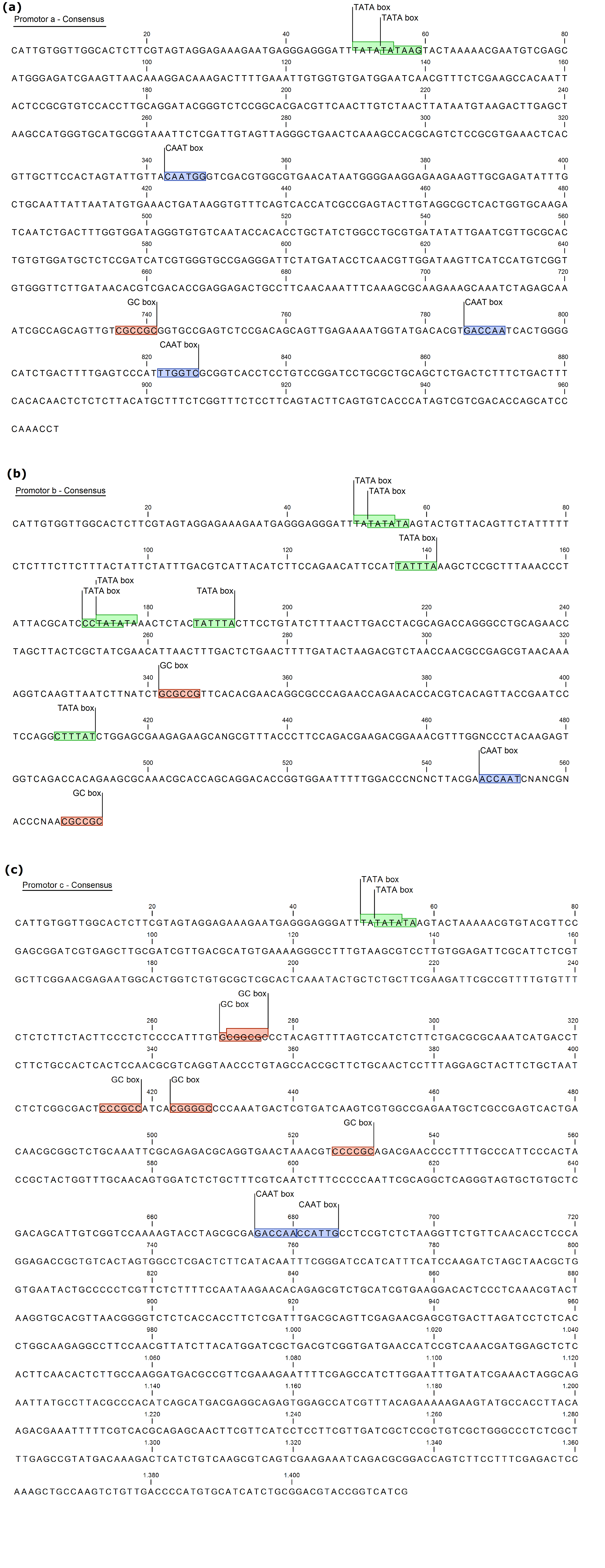
