## Supplemental method S1 for "Adaptive zinc tolerance is supported by extensive gene multiplication and differences in cis-regulation of a CDF transporter in an ectomycorrhizal fungus"

***DNA extraction (modified from Liao et al. 2014)***

Fungal tissue was ground with mortar and pestle until a very fine powder was formed. Cells were lysed with 2% cetyltrimethylammonium bromide (CTAB) lysis buffer Biochemica (ITW Reagents, Darmstadt, Germany) and incubated for 1h at 65°C. Subsequently, DNA was phenol/chloroform extracted. After a first purification step with phenol:chloroform:isoamyl alcohol (25:24:1), the resulting aqueous phase was incubated at room temperature for 20min with guanidinium hydrochloride extraction (GHCL) buffer (6.5M guanidium hydrochloride,100mM Tris-HCL, 0.1M sodium acetate (NaOAc), 0.2M potassium acetate (KOAc), 0.1M β-mercaptoethanol) (Dos Reis et al., 2008). A second and third purification step were carried out with phenol:chloroform:isoamyl alcohol (25:24:1) and chloroform:isoamyl alcohol (24:1), respectively. Ethanol (EtOH) 99% and 3M NaOAc were added to the aqueous phase and samples were incubated overnight at -20°C. After incubation, two 70% EtOH precipitations and one 99% EtOH precipitation were performed. The resulting pellet was resuspended in nuclease-free $H_2O$ and samples were ribonuclease (RNase) treated with RNase Cocktail™ Enzyme Mix (Thermo Fisher Scientific, Waltham, Massachusetts, US) and incubated for 3h at 37°C. After RNase treatment another 99% EtOH precipitation was performed and the DNA pellet was dissolved in nuclease-free $H_2O$.
