## Supplemental method S2 for "Adaptive zinc tolerance is supported by extensive gene multiplication and differences in cis-regulation of a CDF transporter in an ectomycorrhizal fungus"

***PCR SlZnT2 promotor genotypes***

Primers for the PCR amplification of the *SlZnT2* promotor genotypes were designed with Primer3web version 4.1.0 software (Rozen and Skaletsky, 2000) and are listed in Table S2. Each reverse primer is located on the *SlZnT2* gene.

The Platinum Taq DNA polymerase high fidelity kit (Invitrogen, Carlsbad, California, US) was used according to the manufacturer's instructions. 0.5ng DNA was used as input for each PCR reaction. PCRs had a total reaction volume of 50µl.

PCR cycling conditions are specified below in Table M1.

**Table M1** PCR *SlZnT2* promotor genotypes cycling conditions.

| **PCR promotor a** | | | |
|---|---|---|---|
| **Step** | | **Temperature (°C)** | **Time (s)** |
| Initial Denaturation | | 94 | 120 |
| 35 PCR cycles | Denature | 94 | 15 |
| | Anneal | 55.4 | 30 |
| | Extend | 68 | 80 |
| Hold | | 4 | ∞ |
| **PCR promotor b** | | | |
| **Step** | | **Temperature (°C)** | **Time** |
| Initial Denaturation | | 94 | 120 |
| 35 PCR cycles | Denature | 94 | 15 |
| | Anneal | 55.6 | 30 |
| | Extend | 68 | 80 |
| Hold | | 4 | ∞ |
| **PCR promotor c** | | | |
| **Step** | | **Temperature (°C)** | **Time** |
| Initial Denaturation | | 94 | 120 |
| 35 PCR cycles | Denature | 94 | 15 |
| | Anneal | 56.1 | 30 |
| | Extend | 68 | 100 |
| Hold | | 4 | ∞ |
