## Supplementary material for "Adaptive zinc tolerance is supported by extensive gene multiplication and differences in cis-regulation of a CDF transporter in an ectomycorrhizal fungus": Table S1

Table S1: Primer sequences of primers used for gene expression measurements, copy number assessment, and *SlZnT2* promotor amplification.

| **Primers gene expression (RT-qPCR)** | |
|---|---|
| **Primer Name** | **Primer sequence (5'→ 3')** |
| ZnT1F<br>ZnT1R | GGTACTCGGGTTGAATAGTACTAGAATGT<br>GGCATCACGAAACCATGACTTCAT |
| ZnT2F<br>ZnT2R | CGACGGTAAGGTGGAAATAAAC<br>TGGTGAGCCAGATGACAAGA |
| MTaF<br>MTaR | TGTGTCAACAGCGCACATTC<br>GCACCCATTCATCATCACTTT |
| ZRT1F<br>ZRT1R | GCCAAACGGACAAACTGG<br>GACAGGCACGGAGATGAAAG |
| GSS_F<br>GSS_R | CTTTGGGTATGCGTTGTTTG<br>CCGCCCTCATTACTCTCTGT |
| GST_F<br>GST_R | CTTTGAGGGCGAGGATGGATTC<br>TGTTTCCAAGAAGCCCAGACTCAG |
| MnSOD_F<br>MnSOD_R | ATCCGATCGTCACGACTGC<br>TTAAGGTACTGGAGGTAGAAAGCA |
| HSP70F<br>HSP70R | CATCTTGGCATCCTTGAGTACCTT<br>AGGACTTCTCTGCCAACATCAC |
| **Primers gene copy number assessment (qPCR)** | |
| **Primer Name** | **Primer sequence (5'→ 3')** |
| ACT1_1F[1]<br>ACT1_1R[1] | CTGCCTTGTGGTCTGAGTTT<br>GACTGGTGAGAGATGGAGGTT |
| ACT1_2F[1]<br>ACT1_2R[1] | CGCCTGAGCGGAAATACTC<br>CATACTGCGATGAACGATACCA |
| SlZnT2_1F[1]<br>SlZnT2_1R[1] | AATCCCTCCCTCATTCTTTCTC<br>TCACCGACGACGAGTTCTAA |
| SlZnT2_2F[1]<br>SlZnT2_2R[1] | ATCCATTCCAGCACTATTCAGC<br>GAATCAAGCAAGGAGAATCCAC |
| [1] all primer pair efficiencies were between 85 - 115% | |
| **Primers for *SlZnT2* promotor region amplification** | |
| **Primer Name** | **Primer sequence (5'→ 3')** |
| SlZnT2_prom1F[2]<br>SlZnT2_prom1R[2] | GCAAAAGAGGCGTAAGGAGTT<br>AGCCGTAGGAGTAACGAGAATG |
| SlZnT2_prom2F[2]<br>SlZnT2_prom2R[2] | CCNATTTCTTTAGGCATGTCGT<br>GGCGAGGTAGTTGAAATAGAGC |
| SlZnT2_prom3F[2]<br>SlZnT2_prom3R[2] | TGTGTTCCTTTCGTCGAAGTCT<br>AGAGCAAGAGAACCAACGACAT |
| [2] prom1 = promotor genotype 1, prom 2 = promotor genotype 2, and prom 3 = promoter genotype 3. | |

Table S2: Genes of interest description and JGI Protein ID numbers.

| Candidate genes | | |
|---|---|---|
| **Name** | **JGI protein ID**[1] | **Description** |
| *SlZnT1* | 2846331 | Vacuolar Zn transporter of the CDF family (Ruytinx et al., 2017) |
| *SlZnT2* | 2854961 | Zn transporter of the CDF family (Ruytinx et al., 2017) |
| *SlMTa* | 2815822 | Cu metallothionein (Nguyen et al., 2017) |
| *SlZRT1* | 2764984 | Plasma membrane-located ZIP transporter (Coninx et al., 2017) |
| *HSP70* | 2898938 | Molecular chaperones HSP70/HSC70, HSP70 superfamily (KOG Description)[2] |
| *MnSOD* | 2470845 | Manganese superoxide dismutase (KOG Description)[2] |
| *GST* | 2787626 | Glutathione S-transferase (KOG Description)[2] |
| *GSS* | 2771064 | Glutathione synthetase (KOG Description)[2] |

[1] *S. luteus* reference genome UH-Slu-Lm8-n1 v3.0 (Kohler et al., 2015)

[2] KOG = EuKaryotic Orthologous Groups

Table S3: Overview of all the *S. luteus* isolates included in this study together with their geographical collection sites. All collection sites were located in the province Limburg (Belgium). Characteristics of the isolates and collection sites were described previously (Colpaert et al., 2004; Muller et al., 2004).

| **Isolate** | **Abbreviation** | **Geographical collection sites in the province Limburg (Belgium)** |
|---|---|---|
| UH_Slu_DS3 | DS3 | Dilsen-Stokkem |
| UH_Slu_EW11 | EW11 | Eksel |
| UH_Slu_EW12 | EW12 | Eksel |
| UH_Slu_EW2 | EW2 | Eksel |
| UH_Slu_EW4 | EW4 | Eksel |
| UH_Slu_EW5 | EW5 | Eksel |
| UH_Slu_HH1 | HH1 | Hechtelse heide |
| UH_Slu_HH10 | HH10 | Hechtelse heide |
| UH_Slu_HH17 | HH17 | Hechtelse heide |
| UH_Slu_HH18 | HH18 | Hechtelse heide |
| UH_Slu_HH19 | HH19 | Hechtelse heide |
| UH_Slu_HH5 | HH5 | Hechtelse heide |
| UH_Slu_HR1 | HR1 | Houthalen remo |
| UH_Slu_LC2 | LC2 | Lommel containerpark |
| UH_Slu_LM2 | LM2 | Lommel Maatheide |
| UH_Slu_LM3 | LM3 | Lommel Maatheide |
| UH_Slu_LM5 | LM5 | Lommel Maatheide |
| UH_Slu_LM8 | LM8 | Lommel Maatheide |
| UH_Slu_LS1 | LS1 | Lommel sahara |
| UH_Slu_LS4 | LS4 | Lommel sahara |
| UH_Slu_LS8 | LS8 | Lommel sahara |
| UH_Slu_MG4 | MG4 | Meeuwen-Gruitrode |
| UH_Slu_MM4 | MM4 | Maasmechelen |
| UH_Slu_MM7 | MM7 | Maasmechelen |
| UH_Slu_N3 | N3 | Neerpelt |
| UH_Slu_N4 | N4 | Neerpelt |
| UH_Slu_N5 | N5 | Neerpelt |
| UH_Slu_OF3 | OF3 | Overpelt fabriek |
| UH_Slu_OF6 | OF6 | Overpelt fabriek |
| UH_Slu_OF8 | OF8 | Overpelt fabriek |
| UH_Slu_P1 | P1 | Paal |
| UH_Slu_P13 | P13 | Paal |
| UH_Slu_P14 | P14 | Paal |
| UH_Slu_P2 | P2 | Paal |
| UH_Slu_P4 | P4 | Paal |
| UH_Slu_P8 | P8 | Paal |
